# Nucleoside reverse transcriptase inhibitors are the major class of HIV antiretroviral therapeutics that induce neuropathic pain in mice

**DOI:** 10.1101/2022.01.27.478061

**Authors:** Keegan Bush, Yogesh Wairkar, Shao-Jun Tang

**Author notes:** Corresponding Authors Corresponding to: Shao-Jun Tang,; Yogesh Wairkar.

## Abstract

The development of combination antiretroviral therapy (cART) has transformed human immunodeficiency virus (HIV) infection from a lethal diagnosis into a chronic disease, and people living with HIV on cART can experience an almost normal life expectancy. However, these individuals often develop various complications that lead to decreased quality of life, one of the most significant of which is neuropathic pain and development of painful peripheral sensory neuropathy (PSN). Critically, although cART is thought to induce pain pathogenesis, the relative contribution of different classes of antiretrovirals has not been systematically investigated. In this study, we measured development of pathological pain and peripheral neuropathy in mice orally treated with distinct antiretrovirals at their translational dosages. Our results show that only nucleoside reverse transcriptases (NRTIs), but not other types of antiretrovirals, such as proteinase inhibitors, non-nucleoside reverse transcriptase inhibitors, integrase strand transfer inhibitors, and CCR5 antagonists, induce pathological pain and PSN. Thus, these findings suggest that NRTIs are the major class of antiretrovirals in cART that promote development of neuropathic pain. As NRTIs form the essential backbone of multiple different current cART regimens, it is of paramount clinical importance to better understand the underlying mechanism to facilitate design of less toxic forms of these drugs and or potential mitigation strategies.

## Introduction

Antiretrovirals targeting human immunodeficiency virus (HIV) have revolutionized treatment for this deadly disease, and these drugs continue to be an active avenue of investigation, both to identify new compounds that inhibit viral replication and to better understand the secondary symptoms they induce in patients. Combination antiretroviral therapy (cART) effectively inhibits HIV replication in patients, drastically increasing life expectancy post-infection^1^. However, although the profile of HIV-associated neurocognitive disorder (HAND) has shifted away from dementia and severe neurocognitive disorder in the cART era^2^, the prevalence of peripheral sensory neuropathy (PSN) has remained unchanged at around 30–60%, with the variation likely arising from different assessment methods and variable inclusion of decreased sensation neuropathy^3–5^. The leading hypothesis to explain these trends proposes that cART is involved in the initiation and perpetuation of neuropathic symptoms^2,6–11^.

Many studies have demonstrated the potential for several cART components to inhibit neuronal function and induce neurodegeneration. However, these neurotoxic effects are highly dependent on antiretroviral classification^12–25,25–36^. In particular, nucleoside reverse transcriptase inhibitors (NRTIs) are the primary component of cART implicated in PSN development^12,36–39^. These were the first class of antiretrovirals to be used for HIV^10^ and remain one of the most efficacious drugs for long-term treatment of HIV-positive patients. Thus, despite their known detrimental effects, use of NRTIs in patients cannot be discontinued.

Other classes of cART therapeutics include non-nucleoside reverse transcriptase inhibitors (NNRTIs), viral entry inhibitors, which can be separated into C-C chemokine receptor type 5 antagonists (CCR5As) and fusion inhibitors (FIs), integrase strand transfer inhibitors (INSTIs), sometimes referred to integrase inhibitors (IIs), and protease inhibitors (PIs). In general, NNRTIs are one of the most well-tolerated components of cART, with few side effects or drug–drug interactions^9,10,40^. One exception, however, is efavirenz, which has been shown to induce sleep disorders and detrimental psychiatric symptoms in patients and to promote neurodegenerative effects *in vitro* and *in vivo*^20,31–34^. In addition, NNRTIs display one of the lower barriers to resistance among cART components^41,42^. Viral entry inhibitors also tend to be well tolerated, although they carry a slight risk of hepatotoxicity^13,43,44^. However, the injectable nature of FIs has limited their use. This is especially true of enfuvirtide, which requires *bis in die* (twice per day) injection. In contrast, ibalizumab, a newer FI, only requires injection once per week or even once every two weeks and therefore may be more accessible to patients^14–16,45–47^. INSTIs are the second most utilized cART component, often prescribed with two NRTIs for an initial cART regimen^17,48–51^. INSTIs also tend to be relatively well tolerated, with some evidence linking them to insomnia in patients^18,19^. On the other hand, PIs display a variable array of off-target effects, with evidence linking these drugs to development of insulin resistance, increased risk of heart disease, lipodystrophy, and hepatotoxicity, as well as a high potential for drug–drug interaction^23,28,52–56^. Notably, although there is no direct evidence for PIs inducing PSN at clinically relevant levels, some have speculated they may promote exacerbation of NRTI-induced neuropathy^26,27^.

In numerous studies, these various drug classes have been investigated *in vivo* and *in vitro* to determine and confirm their effects, many of which have recapitulated observed outcomes in patients. However, a number of animal models used to test the nociceptive effects of various cART components have utilized injection as the medium for administration. For FIs, this dosing method mimics patient administration; however, NRTIs, NNRTIs, CCR5As, and PIs are administered as ingested pills. Thus, the use of injection administration in these animal models excludes the effects of first-pass metabolism, and critically, several lines of evidence suggest that secondary metabolites of these drugs are involved in their secondary effects^10,23,33,52,57^. Therefore, we hypothesize that administering the commonly ingested cART components to mice via the oral route will more closely mimic the observed effects in patients and lead to the induction of nociception and peripheral neurodegeneration exclusively with NRTI drugs.

Here, to test this hypothesis, we developed a water-based ingestion regime for mice at the translated equivalent dose of each drug class, which was based on methods from past studies utilizing corresponding water-treatment methods^58–60^. We measured the thermal and mechanical nociception profile of each mouse throughout the 4-week dosing regimen, followed by sacrifice and tissue acquisition for further processing. Consistent with our hypothesis, we found that treatment with NRTI induces an increase in nociceptive sensitivity, as well as a decrease in epidermal innervation of the hind paw. These neuropathic phenotypes mimic patient PSN presentation, suggesting that drug-induced neuropathy translates well in our mouse model^3,4,61^. In contrast, the representative NNRTI, CCR5A, and PI drugs did not induce nociceptive or epidermal innervation changes compared to control. Notably, the close recapitulation of observed patient responses to these compounds observed in this study suggests that our ingested administration model may be utilized to further investigate the secondary effects of cART components in future studies.

## Materials and Methods

### Animals

These experiments were performed with adult C57BL/6J mice (10–18 weeks old and weighing 20–32 g) purchased from The Jackson Laboratory (Bar Harbor, ME, USA). All experimental procedures were approved by the Institutional Animal Care and Use Committee at the University of Texas Medical Branch (Protocol #1804026). Thermal and mechanical nociception testing was performed following the guidelines of the International Association for the Study of Pain. Mice were housed in cages (≤5 animals/cage) with standard bedding and free access to food and water, in a room maintained at 23±3°C and a 12/12 light–dark cycle.

### Mouse thermal nociception

Mouse thermal nociception was assessed by tail flick testing, as described previously^62,63^. For these experiments, a water bath was heated to 48°C with precise monitoring of temperature, as even temperature fluctuations of 1–2°C may significantly affect mouse response time. Mice were placed in plastic restraining cones with small open ends (smaller than the standard nose opening size) and allowed to adjust themselves until their tail protruded from the small cone end. Mice will normally self-insert their tails through the small opening; however, if they had trouble doing so, we assisted by either angling the cone slightly or retrying after slight enlargement of the cone tip opening. The larger cone section was then gently folded over the body and head of the mouse to restrain it. Note that restraining cones were modified to have nose holes at the area of the head of the mouse for nose protrusion during the restrained steps. Mice were then positioned above the hot water bath, and a timer was started at the point of tail immersion. The timer was stopped upon tail twitch/flick or retraction, and the time was recorded. Note that in some instances, mice reacted prior to full tail immersion. In this case, we removed the tail from the water bath, let it rest in the air for 30 s, and retried. If tail flick was again observed prior to full tail submersion, time was recorded as 0 s.

### Mouse mechanical nociception

Mouse mechanical nociception was assessed by von Frey testing, as described previously^36,62^. Mice were habituated in 13.2×5×4-cm Plexiglas boxes for 1 h on three continuous days prior to behavioral testing. On testing days, mice were first put in boxes for 20 min to minimize their exploring behavior. The test was then performed by applying the tip of a 3.61/0.4-g filament (Stoelting, Wood Dale, IL, USA) perpendicularly to the center plantar area of the mouse hind paw until the filament was slightly bent (roughly 30° from tip to tip) and remained bent for one second. Mouse response and filament switching was based on the up–down method. That is, if the mouse responded to the initial filament, we utilized the next smaller force filament, and if there was no response, we utilized the next larger force filament. This process was repeated until six up–down measurements were acquired per paw per mouse at each measurement time. Nociception withdraw response was recorded for abrupt paw withdrawal, lateral moving, shaking, lifting, licking, and/or biting of the touched area of the hind paw.

### Mouse treatment

Mouse treatment with cART was administered via water ingestion^58–60^. Each drug was thoroughly dissolved in an initial water volume of 100 mL, which was provided to the mice and refilled every day, or every other day (depending on the observed fill line), with freshly made 50-mL aliquots of the same drug, at the same relative concentration. Water bottles in cages were closely monitored daily for ingestion and any precipitation of drug; all drugs are readily soluble at the administered doses, and no precipitation was observed. Concentrations for each cART component in water were as follows: 0.098-mg/ml emtricitabine (FTC; NRTI), 0.59-mg/ml ritonavir (PI), 0.39-mg/ml raltegravir (II/INSTI), 0.29-mg/ml efavirenz (NNRTI), and 0.29-mg/ml maraviroc (CCR5A); control mice received water without drug. At the average water consumption of 5-mL/day for mice of this age and weight, these concentrations roughly equate to the human equivalent of these drugs at their most common dosage regimens in patients. Water level changes were also monitored to confirm ingestion amounts. We detected no changes in water consumption within groups pre- and post-initiation of treatment or between the different treatment groups.

### Mouse dissection, imaging, and analysis

Dissection of the hind paw skin was performed as described previously^64,65^. Briefly, mice were anesthetized with 14% urethane (0.2–0.3 ml; intraperitoneally) and monitored for complete anesthetization via hind paw pinch prior to decapitation. After other dissection tissues were stored, hind paw skin was collected from both paws and placed in ice-cold 80% ethanol in phosphate-buffered saline (PBS) overnight (or until further processing). Skin sections were flat mounted on ice-cold gel plates and rehydrated in a series of ethanol solutions (50% EtOH, 30% EtOH, and 0% EtOH in PBS) prior to cryosectioning (40 μm) and staining with rabbit anti-PGP9.5. Neuronal innervation was quantified by manual counting of epidermal fibers in 100-μm-length segments from 10 random sections located in a similar plantar area of each mouse; sections from both hind paws of individual mice were grouped together.

### Statistical analysis

Statistical analyses were performed using GraphPad Prism software. Significance of skin innervation data comprising single data points per mouse were analyzed using the Student’s *t*-test for comparison between two groups and one-way analysis of variance (ANOVA), followed by the Bonferroni posthoc test, for comparisons between multiple groups. Significance of results from analyses yielding multiple data points per mouse (von Fray and tail flick data) was determined by two-way ANOVA, followed by the Bonferroni multiple comparisons test. Significance levels were set to 95% confidence intervals, and *P*-values are indicated in the respective figure legends.

## Results

### cART induces mechanical nociception changes

Previous studies using mammalian animal models have reported mechanical nociception changes in response to NRTI and PI admiminstration^27,36,53,66–68^. However, in many of these investigations, the drugs were administered via injection, and several of the NRTI, and all of the PI, studies that detected nociception changes used dosing regimens that translate above clinically relevant levels. Thus, although these studies have reported neurodegenerative effects, they do not mimic patient administration conditions. Here, utilizing a mouse model with clinically relevant translated doses and oral drug administration, we hypothesized that mice treated with an NRTI would exhibit a decreased mechanical nociceptive threshold, whereas in those administered a PI, II, CCR5 antagonist, or NNRTI, the exhibited mechanical nociception would remain constant.

To test this hypothesis, we utilized a previously developed model for azidothymidine (AZT) administration via the water route as a framework for our dosing regimens. This study reported that administration of AZT causes no visible distress to the animals and no changes in water intake, and the drug shows stability in water^58–60^. Using the known translation index for mice by weight, combined with the average water intake of mice, we then translated the most commonly administered patient dose for each cART component. For the NRTI and CCR5 antagonist, we administered the most commonly prescribed variant, whereas for the II, PI, and NNRTI, we administered the therapeutic variant that exhibits the most neurotoxic potential. This decision was made based on the notably enhanced neurodegenerative potential demonstrated by one form of these drugs relative to others in the same class^18,20,31–34,52,53,57^ and our goal of testing PSN-inducing ability of the worst-case patient therapeutic regimen for each cART component. In this model, mice ingested individual cART components over a period of 4 weeks and were monitored for mechanical nociceptive changes throughout their treatment via von Frey filament testing. Consistent with our predictions, we detected no deviation in mechanical nociception throughout the 4-week treatment regimen in mice administered PI, II, CCR5 antagonist, or NNRTI relative to controls (Figs. 2–5A). In contrast, mice treated with NRTI (FTC) exhibit a progressive decrease in mechanical nociceptive threshold over a period of 3 weeks and a near steady allodynia-like phenotype for the last week of treatment (Fig. 1A). This suggests that NRTIs show greater potential for induction of mechanical sensitization when given under conditions that closely mimic those for patient administration.

**Figure 1:**
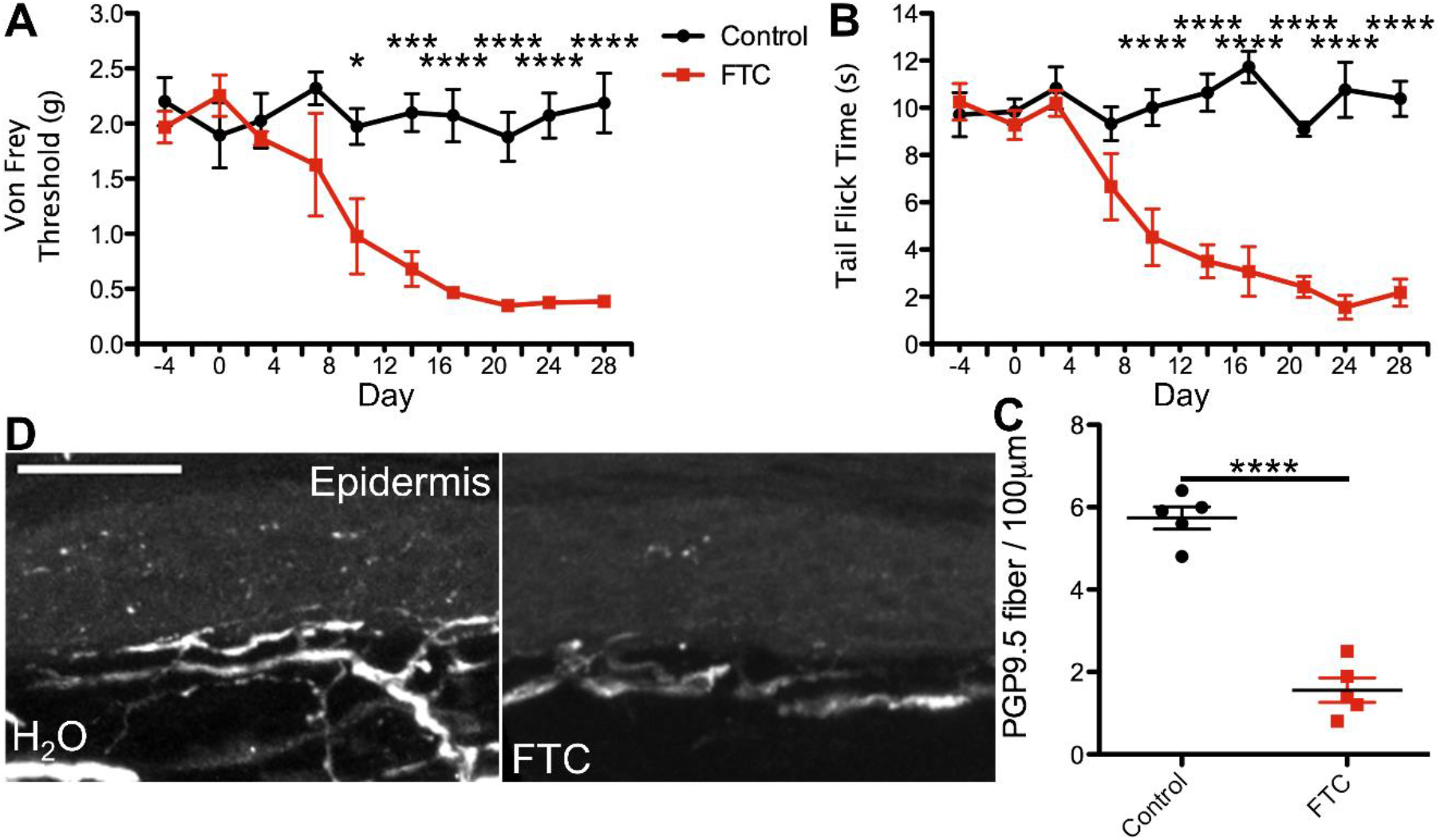
Nucleoside reverse transcriptase inhibitor (NRTI) orally administered at the translated patient dosage induces nociception sensitization and epidermal denervation in mice. **A**) Average von Frey filament threshold in grams throughout the 4-week treatment regimen for mice administered 0.098-mg/mL emtricitabine (FTC; concentration used in all experiments), a commonly used NRTI, in water and control animals (*n* = 5, both groups). **B**) Average tail flick times in seconds throughout the 4-week treatment regimen for FTC-treated and control mice. **C**) Number of PGP9.5-labeled axons per 100 μm, averaged from 10 sections per mouse, in FTC-treated and control mice. **D**) Representative images of PGP9.5-labeled axons in hind paw glabrous skin epidermis from FTC-treated and control mice. \**P* <0.05, \*\*\**P* <0.001, \*\*\*\**P* <0.0001; error bars show the standard error of the mean (S.E.M.); scale bar, 50 μm.

### Thermal nociception sensitization induced by cART

HIV patients with PSN report an array of pain conditions, including numbness, increased sensitivity to touch or thermal extremes, and even, in some cases, chronic pain^3,4,69^. These varied pain profiles likely result from a number of factors; however, reports of PSN linked to NRTI tend to involve mechanical and thermal components. Therefore, we hypothesized that changes in thermal nociception occurring in response to ingestion of individual cART components in our model would mirror any mechanical nociception changes.

To test this possibility, we monitored all mice for any alterations in thermal nociception throughout the 4-week treatment regimen, using the hot water bath tail flick test^62,63^. This method of thermal nociception assessment was chosen over the more commonly used hot plate test due to our need for continued use, as repeated hot plate testing may induce tissue damage and skew the results. In addition, this method of thermal nociception assessment utilizes a different appendage than the hind paw, which is used in von Frey testing, potentially providing a more generalized insight into the mouse nociception profile in response to cART. As hypothesized, we observed no changes in thermal nociception over the 4-week dosing regimen in mice administered PI, II, CCR5A, or NNRTI relative to control animals (Figs. 2–5B). Conversely, mice administered NRTI (FTC) display faster response to a fixed temperature water bath, suggesting an increased thermal sensitization that occurs in parallel with the changes in mechanical nociception described above (Fig. 1B). Thus, these data indicate that the cART component implicated in development of mechanical nociception sensitization also induces changes in thermal nociception in an ingestion mouse model.

**Figure 2:**
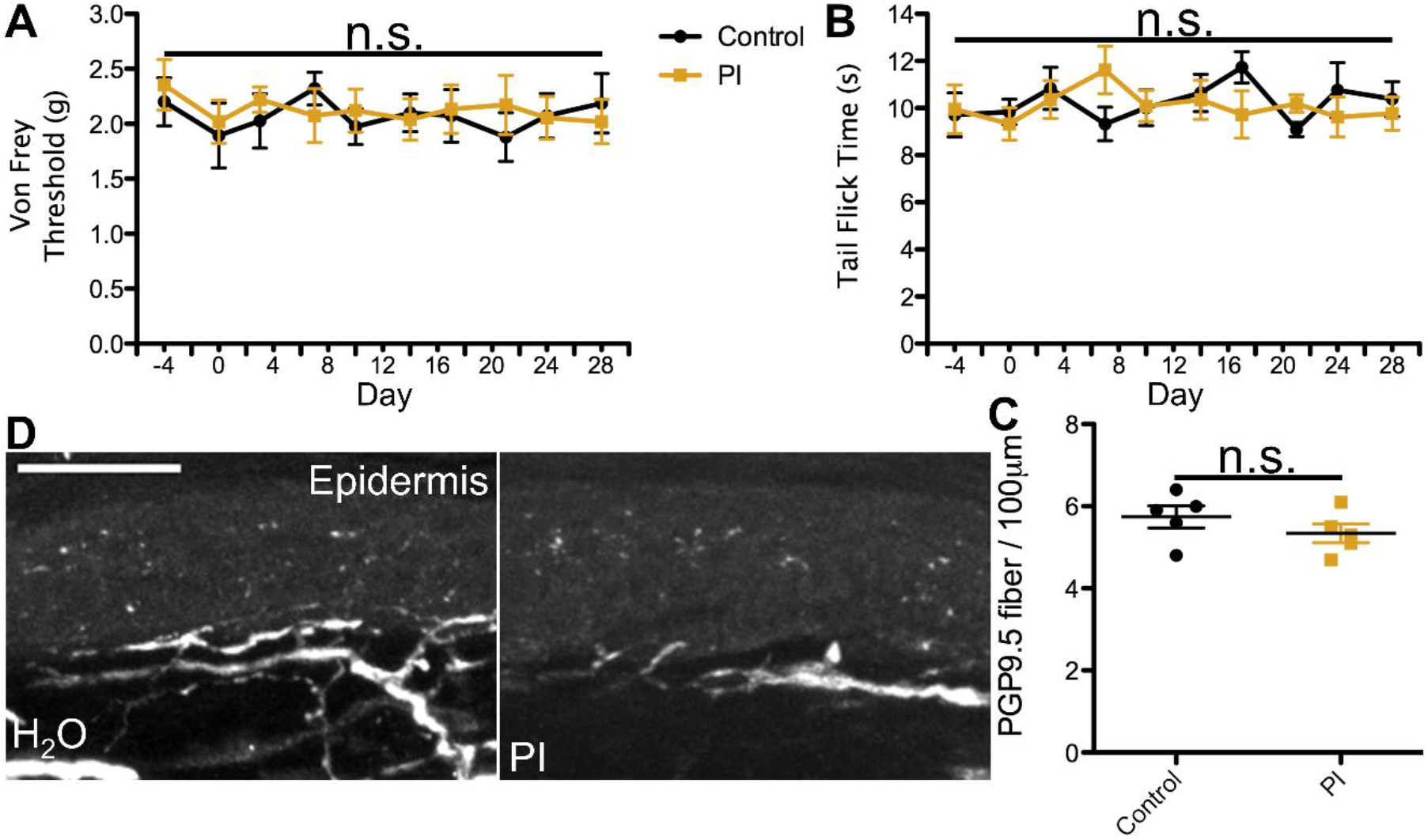
Protease inhibitor (PI) orally administered at the translated patient dosage does not induce changes in nociception or epidermal innervation in mice. **A**) Average von Frey filament threshold in grams throughout the 4-week treatment regimen for mice administered 0.59-mg/ml ritonavir (PI; concentration used in all experiments) in water and control animals (*n* = 5, both groups). **B**) Average tail flick times in seconds throughout the 4-week treatment regimen for ritonavir-treated and control mice. **C**) Number of PGP9.5-labeled axons per 100 μm, averaged from 10 sections per mouse, in ritonavir-treated and control mice. **D**) Representative images of PGP9.5-labeled axons in hind paw glabrous skin epidermis from ritonavir-treated and control mice. No significant differences in nociception or epidermal innervation relative to control were observed at any time point. Error bars show the S.E.M.; scale bar, 50 μm.

**Figure 3:**
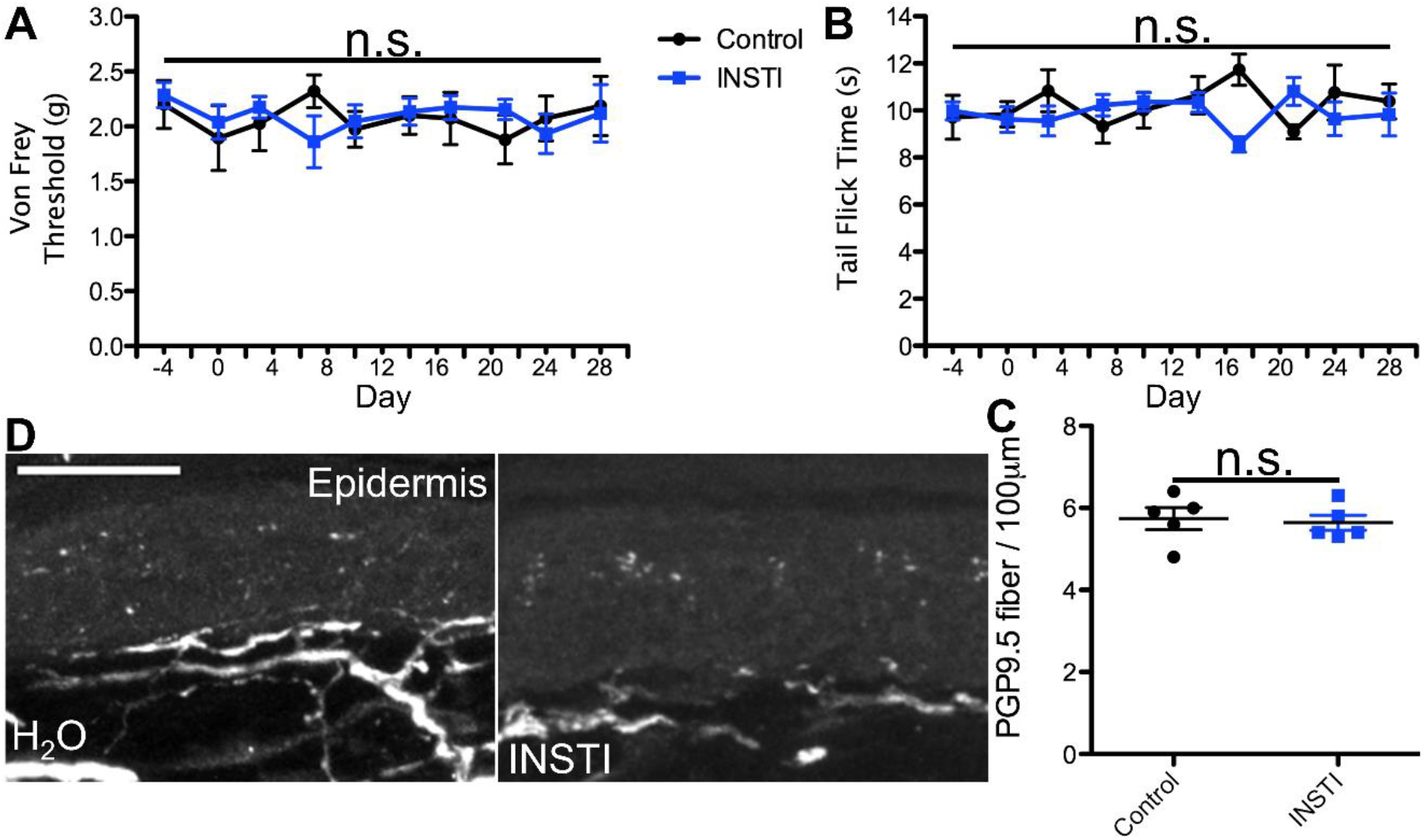
Integrase strand transfer inhibitor (INSTI) orally administered at the translated patient dosage does not induce changes in nociception or epidermal innervation in mice. **A**) Average von Frey filament threshold in grams throughout the 4-week treatment regimen for mice administered 0.39-mg/ml raltegravir (INSTI; concentration used in all experiments) in water and control animals (*n* = 5, both groups). **B**) Average tail flick times in seconds throughout the 4-week treatment regimen for raltegravir-treated and control mice. **C**) Number of PGP9.5-labeled axons per 100 μm, averaged from 10 sections per mouse, in raltegravir-treated and control mice. **D**) Representative images of PGP9.5-labeled axons in hind paw glabrous skin epidermis from raltegravir-treated and control mice. No significant differences in nociception or epidermal innervation relative to control were observed at any time point. Error bars show the S.E.M.; scale bar, 50 μm.

**Figure 4:**
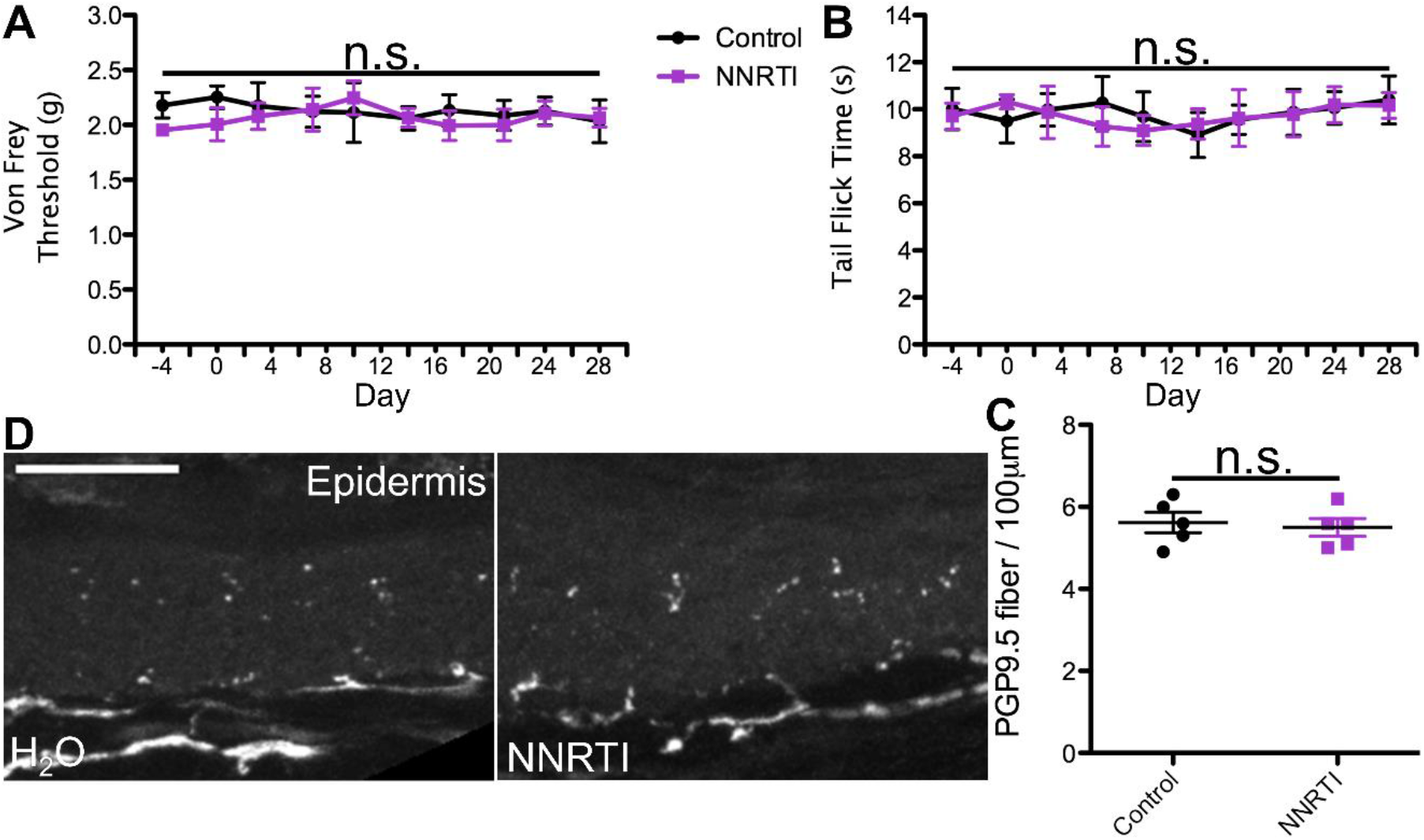
Non-nucleoside reverse transcriptase inhibitor (NNRTI) orally administered at the translated patient dosage does not induce changes in nociception or epidermal innervation in mice. **A**) Average von Frey filament threshold in grams throughout the 4-week treatment regimen for mice administered 0.29-mg/ml efavirenz (NNRTI; concentration used in all experiments) in water and control animals (*n* = 5, both groups). **B**) Average tail flick times in seconds throughout the 4-week treatment regimen for efavirenz-treated and control mice. **C**) Number of PGP9.5-labeled axons per 100 μm, averaged from 10 sections per mouse, in efavirenz-treated and control mice. **D**) Representative images of PGP9.5-labeled axons in hind paw glabrous skin epidermis from efavirenz-treated and control mice. No significant differences in nociception or epidermal innervation relative to control were observed at any time point. Error bars show the S.E.M.; scale bar, 50 μm.

**Figure 5:**
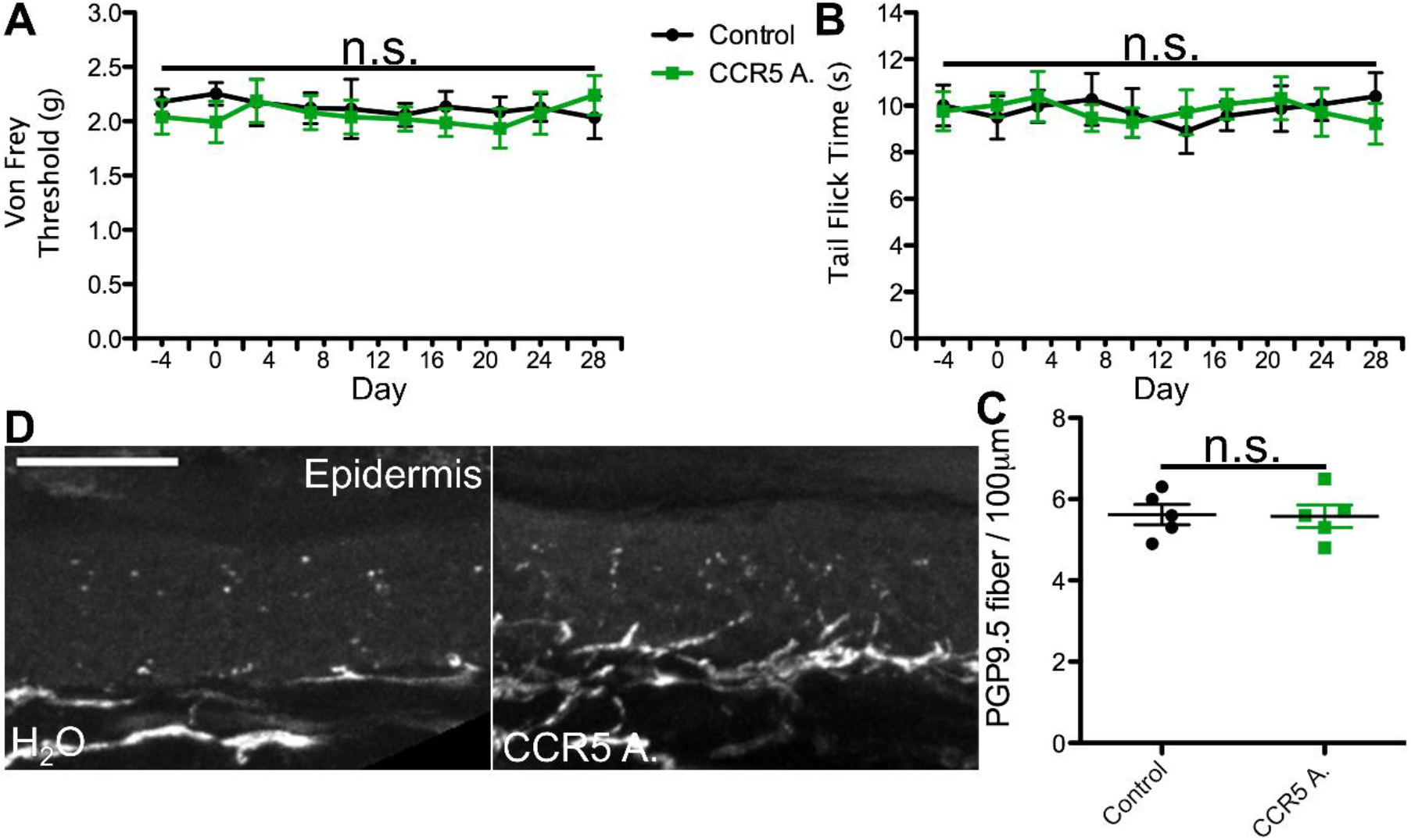
C-C chemokine receptor type 5 antagonist (CCR5A) orally administered at the translated patient dosage does not induce changes in nociception or epidermal innervation in mice. **A**) Average von Frey filament threshold in grams throughout the 4-week treatment regimen for mice administered 0.29-mg/ml maraviroc (CCR5A; concentration used in all experiments) in water and control animals (*n* = 5, both groups). **B**) Average tail flick times in seconds throughout the 4-week treatment regimen for maraviroc-treated and control mice. **C**) Number of PGP9.5-labeled axons per 100 μm, averaged from 10 sections per mouse, in maraviroc-treated and control mice. **D**) Representative images of PGP9.5-labeled axons in hind paw glabrous skin epidermis from maraviroc-treated and control mice. No significant differences in nociception or epidermal innervation relative to control were observed at any time point. Error bars show the S.E.M.; scale bar, 50 μm.

### Epidermal denervation induced by cART

HIV patients with PSN have been examined for changes in epidermal innervation, with neuropathy and epidermal neuron loss observed in some cases^61,70–72^. Therefore, we hypothesized cART components that induce changes in nociception profiles would also promote denervation of the epidermis. To test this possibility, we harvested glabrous hind paw skin tissue at the conclusion of our 4-week cART administration regimen and measured epidermal innervation via PGP9.5 staining. Our data reveal relatively consistent innervation in mice treated with control, II, PI, CCR5A, and NNRTI (Fig. 2–5C,D), whereas we detected a significant decrease in epidermal axon counts in NRTI-treated mice (Fig. 1C,D). This suggests that patterns of epidermal peripheral neuron degeneration occurring in response to oral NRTI in mice may model those observed in PSN patients. Overall, our findings suggest that this ingestion model may show utility for closely modeling and uncovering the mechanism for NRTI-induced PSN-associated pain in patients.

## Discussion

In this study, we measured the effects of individual cART therapeutics, ingested at the translated human dosage equivalent, on nociception and peripheral neurodegeneration in a mouse model. Critically, this water ingestion model more closely mimics the method for patient administration than previous studies that used injection-based administration^27,36,53,58–60,66–68^. We found that when orally administered at clinically relevant levels, PI, II, CCR5A, and NNRTI do not induce detectable changes in nociception and epidermal innervation. In contrast, ingestion of NRTI leads to a maintained thermal and mechanical allodynia-like phenotype, with associated loss of epidermal axon staining. We further note that, to the best of our knowledge, this study is the first to assess nociception and peripheral neurodegeneration in response to ingestion of cART therapeutics in a mouse model.

### Ingestion of cART

In HIV patients, one of the most common cART regimens combines two NRTIs and an INSTI^1,17,48–51^, a formula that is utilized for its efficacy in reducing viral load and the low potential for both drug–drug interactions and development of HIV resistance. However, it is not applicable in all cases, as some individuals exhibit severe acute adverse reactions to this formulation. In such instances, alternative combinations are tested until a well-tolerated and effective regimen is found. Regardless, even for these rare cases, there is a tendency towards the use of therapeutics that are easily administered by patients to promote compliance^14–16,45–47^, which precludes the use of injected cART, if avoidable. Therefore, an animal model that mimics the most common method for patient administration would represent a powerful tool for investigating the effects of cART and the physiological response to these therapeutics. As such, a key aim of our study was to investigate the potential viability and translation relevance of a cART oral administration mouse model.

The inherent difference between ingestion and injection administration primarily derives from two features. The first is the potential effects from first-pass metabolism with ingested drugs, which is avoided in injection. Although the metabolic profiles of many individual cART component variants have not been well characterized, it was shown that PIs are readably metabolized by the CYP3A pathway^52^, which contributes to their drug–drug interactions. In addition, there is evidence that at least one NNRTI metabolite exhibits the potential for increased neurotoxicity *in vitro*^9,10,23,33,52,57,73^. Based on these observations, a model that includes first-pass metabolism of cART components is likely to better recapitulate patient conditions. The second major source of variation between ingestion and injection relates to the relative dynamic pharmacokinetics induced by these distinct forms of drug administration^74–80^. That is, while ingestion may induce a bolus of drug above target dosage absorption, this usually occurs to a much lesser extent relative to injection administration. Thus, injection may show increased potential for effects, such as above target dosages and both off-target and toxic effects compared to ingestion. We therefore expect that our ingestion model has less potential to generate false positives resulting from such effects, again suggesting closer modeling of the patient condition than injection.

### Neurotoxicity of cART

The possible neurotoxicity of many of cART components has been tested *in vitro* and in several animal models. In particular, some PI variants have been shown to induce behavioral deficits, and there is further evidence for PI-induced neurodegeneration in neuron cultures. However, many of these studies used older PI dosing regimens that were known to induce CNS effects in patients, whereas improved protocols for cotreatment with pharmacokinetic enhancers has lowered the effective dosage for PI treatment^27,28,53,81^. Thus, PI concentrations shown to promote neurotoxicity are supraphysiological relative to current patient regimens, and to the best of our knowledge, there have been no studies linking current clinical practice relevant dosages of PI to PSN development^10^. Consistent with this, we detected no nociceptive or epidermal denervation effects from ingested PI at current translated dosages in our mouse model.

IIs are generally considered well tolerated and non-neurotoxic with two exceptions. Raltegravir has been linked to insomnia and anxiety in patients, with *in vitro* studies demonstrating potential for induction of increased stress responses at supraphysiological concentrations^18,20^. Secondly and conversely, studies on dolutegravir are mixed, with some evidence linking it to development of similar neuropsychiatric symptoms, while other evidence suggests that this compound exhibits potential to inhibit neurodegeneration in the CNS^82,83^.

Similar to IIs, most NNRTIs exhibit little to no neurodegenerative effects in patients. The one exception to this is efavirenz, which has been linked to insomnia and development of severe neuropsychiatric disorders. In addition, several animal and *in vitro* studies have demonstrated significant efavirenz-induced neurotoxicity, with evidence for mitochondrial dysfunction^20,31,34^, although we note that these studies did not assess the potential for peripheral neurodegeneration or measure nociception profiles in response to efavirenz.

Similarly, there have been no studies reporting neurotoxic effects from CCR5A administration. In contrast, there is some evidence that intrathecal-injected CCR5A exhibits analgesic effects in a sciatic nerve constriction model, with associated downregulation of proinflammatory markers in microglia^84^.

Distinct from other cART therapeutics, NRTIs are notorious for inducing PSN in patients and in animal models. In particular, there is evidence that several forms of these drugs, including the more recently developed “non-neurotoxic” NRTIs in therapeutic use today, induce painful sensory neuropathy in male and female patients^9,12,35,36,85,86^. However, while NRTI-induced nociception and oral administration have been investigated in animal models individually, to the best of our knowledge, there are no previous reports of nociception changes in response to ingested NRTIs in animals. Therefore, our study is the first to show evidence for nociceptive sensitization and peripheral neurodegeneration induced in response to oral treatment with NRTI in a mouse model.

### Closing Remarks

In this study, we utilized a newly developed mouse ingestion model with individual cART components to assess their nociceptive effects and measure epidermal neurodegeneration in response to these compounds. However, this model has several limitations. The first being that we administered only one component at a time, whereas in patients, cART commonly consists of three or more therapeutics used in conjunction^1,17,48–51^. This difference may lead to discrepancies between the spectrum of conditions exhibited by patients and our model. To address this concern, in future studies, we propose expanding our model to include the assessment of ingested cART combination regimens, in order to investigate the effects of various drug combinations. Secondly, cART is administered in the presence of HIV infection, and the expression of viral products is a potential confounding factor in PSN development^6,87–90^. This can partially be addressed in future studies by testing the effects of oral cART administration in mouse models of HIV gp120 administration or in mice genetically engineered to express viral products^88,90,91^. A third limitation of our study is that an increased prevalence of PSN is observed in older patients^92–94^. This potential risk factor in cART administration can be addressed by performing studies with age matched-mice, which may more closely model this complication. Finally, patient conditions are not static, and often, with modification of cART regimens, PSN symptoms can resolve. Therefore, future investigations focusing on regimen modification of ingested cART may provide additional insight into the interplay between various therapeutics in development and maintenance of PSN presentation. Overall, despite these limitations, this study has yielded a new, broadly applicable ingestion model for cART therapeutics that recapitulates patient administration methods. Critically, this provides a valuable resource for future investigations into the physiological effects of cART treatment and provides a model with which to test possible therapeutic inhibition of cART-induced PSN in response to NRTI or cART administration.

## Notes

### Competing Interest Statement

The authors have declared no competing interest.

